# Modeling wild type and mutant p53 in telomerase-immortalized human cells

**DOI:** 10.1101/2023.06.22.546141

**Authors:** Jessica J. Miciak, Fred Bunz

## Abstract

Genetic alterations that change the functions of p53 or other proteins in the p53 pathway contribute to a majority of cancers. Accordingly, many technological approaches and model systems have been employed to dissect the complex phenotypes of this critical tumor suppressor and its mutants. Studies of human p53 are commonly conducted in tumor-derived cell lines that retain wild type *TP53* alleles and isogenic derivatives with engineered *TP53* alterations. While this genetic approach has provided numerous insights, such studies are bound to paint an incomplete picture of p53 and its many effects on the cell. Given the preponderance of p53 pathway defects in cancer, it is reasonable to assume that cancers that arise without mutations in the *TP53* coding sequence would very likely harbor other genetic or epigenetic alterations that effect the normal function of this pathway. One possible solution to this conundrum is to study p53 in cells that have been artificially immortalized. Unlike cells derived from tumors *ex vivo*, cells that have been immortalized *in vitro* are not shaped by evolutionary selection during tumorigenesis, and presumably retain many of the normal functions of p53 and other tumor suppressors. We report here a functional characterization of p53 in the immortalized human cell line hTERT-RPE1 and describe the dominant-negative effects of a heterozygous missense p53 A276P mutation that apparently arose during serial culture. Detailed studies of this contact mutant, also found in human tumors, demonstrate the practical utility of this model system for studying the complex phenotypes of human p53.

## Introduction

Since its discovery more than four decades ago, p53 has been the subject of intensive investigation. The *TP53* gene is mutated at high frequency in many of the most lethal human cancers [1,2]. A detailed elucidation of p53 function is therefore critical, not only for our understanding of cancer biology, but for the development of therapeutic strategies that preferentially target cells that harbor *TP53* mutations.

Gene editing has been a powerful approach to the study of tumor suppressor pathways in human cells [3]. Despite the high frequency of *TP53* mutations in cancer, many tumor-derived cell lines retain wild type alleles and express wild type p53 protein. Such cell lines, paired with isogenic derivatives that are nullizygous for p53 expression, have found wide use in basic and translational p53 research. Given their utility and accessibility, it is easy to forget that such cell lines cannot be considered normal, nor can their p53 pathways be assumed to be entirely intact. On the contrary, the p53 pathway is so wide-ranging in its cell-autonomous and paracrine functions that one might safely assume that any expanding *TP53*-wild type cell population that passes through the evolutionary bottleneck of tumorigenesis must acquire other stable alterations that allow it to bypass the potent tumor-suppressive effects of p53.

The strengths and limitations of currently available human knockout cell lines are exemplified by the first isogenic cell system for the study of human p53. Generated 25 years ago from the colorectal cancer cell line HCT116 [4] and widely distributed thereafter, the original human p53 knockout cells exhibit many cancer-relevant phenotypes, such as resistance to 5-fluorouracil (5-FU) [5], a first line therapeutic agent for patients with colorectal cancer. This distinctive drug-resistant phenotype, uniquely elicited by 5-FU, established a plausible molecular mechanism for treatment failure. While still widely used for analysis of human p53, the HCT116 model has well-defined limitations. Some of the mechanisms for p53 turnover and induction are reportedly defective in the parental cell line [6]. In addition, HCT116 was derived from a mismatch repair-deficient cancer and harbors >9000 mutations [7]. Most of these alterations are undoubtably passenger mutations of no functional consequence, but the full phenotypic impact of this high mutational burden remains unknown.

A useful system for studying p53 biology in the absence of mutations acquired during tumorigenesis was generated more recently in the breast epithelial cell line MCF-10A [8]. This spontaneously immortalized cell line was derived from apparently normal breast tissue following a mastectomy performed on a premenopausal female with fibrocystic breast disease. Genetically stable and non-tumorigenic, MCF-10A cells are used widely breast cancer research [9]. The biallelic disruption of *TP53* in these cells caused genetic instability and phenotypic heterogeneity in the daughter clones, as well as epidermal growth factor-independent growth. However, in three-dimensional acinar culture, MCF-10A reportedly express ectopic markers of differentiation, suggesting a potential limitation of this system [10]. The underlying mechanisms by which the MCF-10A line initially achieved immortality remain obscure.

To study p53 in a human cell line with a defined basis for immortality, we looked to hTERT-RPE1, hereafter referred to simply as RPE1. This cell line was derived from the primary retinal pigment epithelial cell line RPE-340 [11]. Like other primary cells, RPE-340 cells undergo replicative senescence after 50-60 passages. This limitation, first described by Hayflick, was successfully bypassed by the forced expression of the catalytic subunit of the enzyme telomerase, hTERT. Thus immortalized, RPE1 cells have a stable diploid karyotype, are non-tumorigenic [12] and are widely used as a surrogate for normal cells in studies of cell signaling and cell proliferation.

*In vitro* immortalization by telomerase overexpression has been achieved in diverse human cell lineages. Notably, the bypass of replicative senescence usually requires the introduction of a cellular or viral oncogene, such as SV40 large T antigen, in addition to telomerase. Such oncogenes typically interact with p53 and alter its function. RPE1 cells, immortalized by telomerase expression alone, may therefore be uniquely suited for the study p53 phenotypes in the absence of replicative senescence.

Recent studies employing RPE1 cells and CRISPR-mediated gene editing techniques have demonstrated how p53 suppresses cell growth in response to mitotic dysfunction [13] and maintains genetic stability by preventing polyploidization [14]. Inspired by these elegant studies, we explored the general utility of RPE1 cells for the detailed analysis of the p53 pathway. We found that a diverse array of phenotypes previously attributed to p53, including the induction of numerous downstream transcriptional targets, could be reliably and robustly elicited in RPE1 cells. Many of these phenotypes are absent in other human cell lines that have been similarly assessed. In addition, we describe an unstudied cancer-associated *TP53* mutation that spontaneously arose in this line, and describe how it altered both the transcription-dependent and -independent functions of wild type p53. Our studies suggest that RPE1 is a particularly useful system for studying the basic biology of human p53 and its many mutants

## Results

The *TP53* locus expresses several related proteins from two endogenous promoters (Fig 1A). To eliminate expression of all known p53 isoforms, we transfected RPE1 cells with a CRISPR designed to disrupt exon 7 and, following puromycin selection, isolated individual subclones by limiting dilution. Multiple knockout clones were identified by western blot and confirmed by Sanger sequencing of exon 7.

**Figure 1.**
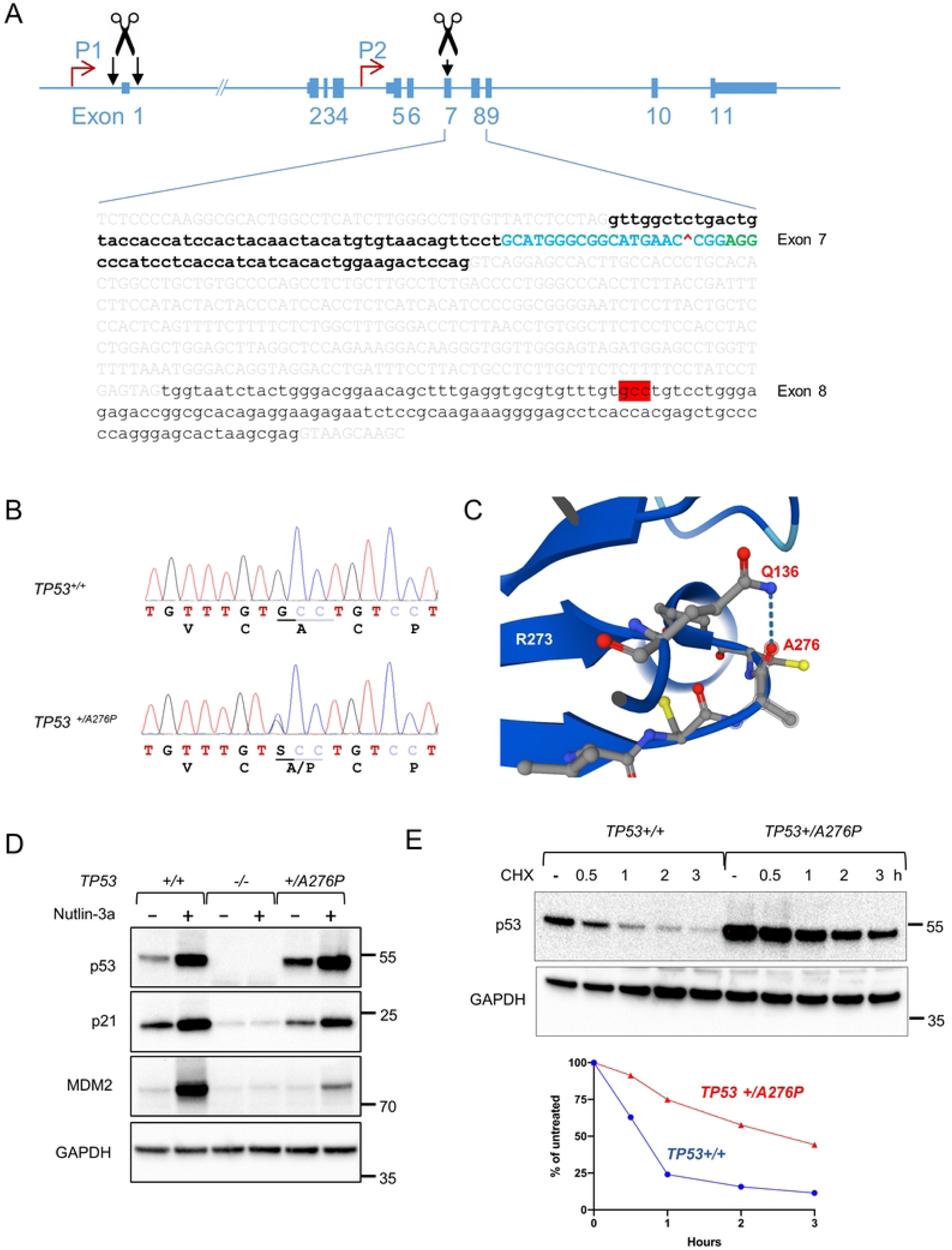
Disruption of the *TP53* locus in RPE1 cells. (A) A guide RNA was designed to target an SpCas9 protospacer sequence in *TP53* exon 7 (shown in blue). The predicted cut site indicated with a red “^” symbol in the text; the protospacer adjacent motif (PAM) is shown in green. Codon 276 is highlighted in red. To create an independent cell line incapable of producing full-length p53, a dual guide approach was used to delete *TP53* exon 1. (B) The Sanger sequence traces indicating the heterozygous A276P mutation, identified in a single clone. (C) A structural analysis predicts a stable hydrogen bond between A276 and Q136. The rendering generated by AlphaFold [82,83] was based on 245 structures available in UniProt P04637. (D) Cells with the indicated genotypes were untreated or treated with 10 µM nutlin-3a for 8 h. Protein extracts were probed for the indicated proteins. (E) Cell were treated with 100 µg/ml cycloheximide for 3 h and harvested at the indicated time points. Protein extracts were probed for p53 and GAPDH (upper panel). Levels of protein were quantified by densitometry (lower panel). The migration of relevant molecular weight markers, in kDa, are shown to the right of each blot.

Unexpectedly, we isolated a single clone that retained wild type exon 7 sequences but expressed elevated levels of p53 protein. As p53 protein overexpression is a consequence of most tumor-associated *TP53* mutations, we decided to investigate further. We sequenced the remaining exons in this clonal cell population and identified a single nucleotide substitution in exon 8 (Fig 1B). A heterozygous C-to-G transversion encoded a proline residue at position 276 in place of the wild type alanine (A276P). This residue is predicted to form a hydrogen bond with Q136 (Fig 1C), and thus may contribute to structural stabilization. According to the TCGA database [15,16], A276P/D/G mutations are found in a relatively small numbers of tumors from diverse tissues and are predicted to be driver mutations.

We expanded the *TP53 +/A276P* cell line to investigate the phenotypic impact of this heterozygous mutation, which is heretofore unstudied. The expression of p53 and two proteins normally induced by p53 were assessed in parental RPE1 and isogenic knockout cells that expressed no p53 protein (Fig 1D). As expected, treatment of RPE1 cells with the MDM2 inhibitor nutlin-3a caused stabilization of p53 and robust induction of p21^WAF1/CIP1^ and MDM2, which are encoded by canonical p53 target genes [17,18]. These proteins were not induced in the knockout cells. As first observed in the initial knockout screen, p53 protein was increased in *TP53 +/A276P* cells and further induced by Nutlin-3a. However, the induction of p21^WAF1/CIP1^ and MDM2 was decreased in these mutant cells compared with wild type cells. A cycloheximide chase experiment (Fig 1E) demonstrated that p53 expressed in the *TP53 +/A276P* cells exhibited increased stability, a cardinal feature of many tumor-associated p53 mutations.

We next sought to determine the effects of wild type p53 and the A276P mutant on cell growth and in the responses to DNA damage. Serially cultured in parallel, the *TP53* -/- and *TP53 +/A276P* cells appeared to reach confluence more rapidly than parental cells, suggesting that p53 might impede cell proliferation in this line. Accordingly, colonies founded by single *TP53* -/- cells were notably larger than colonies derived from wild type cells (Fig 2A). To rule out the possibility of CRISPR off-target effects or simple clonal variation as causes of this growth disparity, we created a deletion in *TP53* exon 1 (Fig 1A) to completely eliminate the expression of full-length p53, but not the remaining isoforms. Notably, the size of the colonies was fairly uniform within each population, suggesting that there was limited variation between subclones. An increased rate of growth in *TP53* -/- and *TP53 +/A276P* cell populations was additionally quantified by time lapse microscopy (Fig 2A).

**Figure 2.**
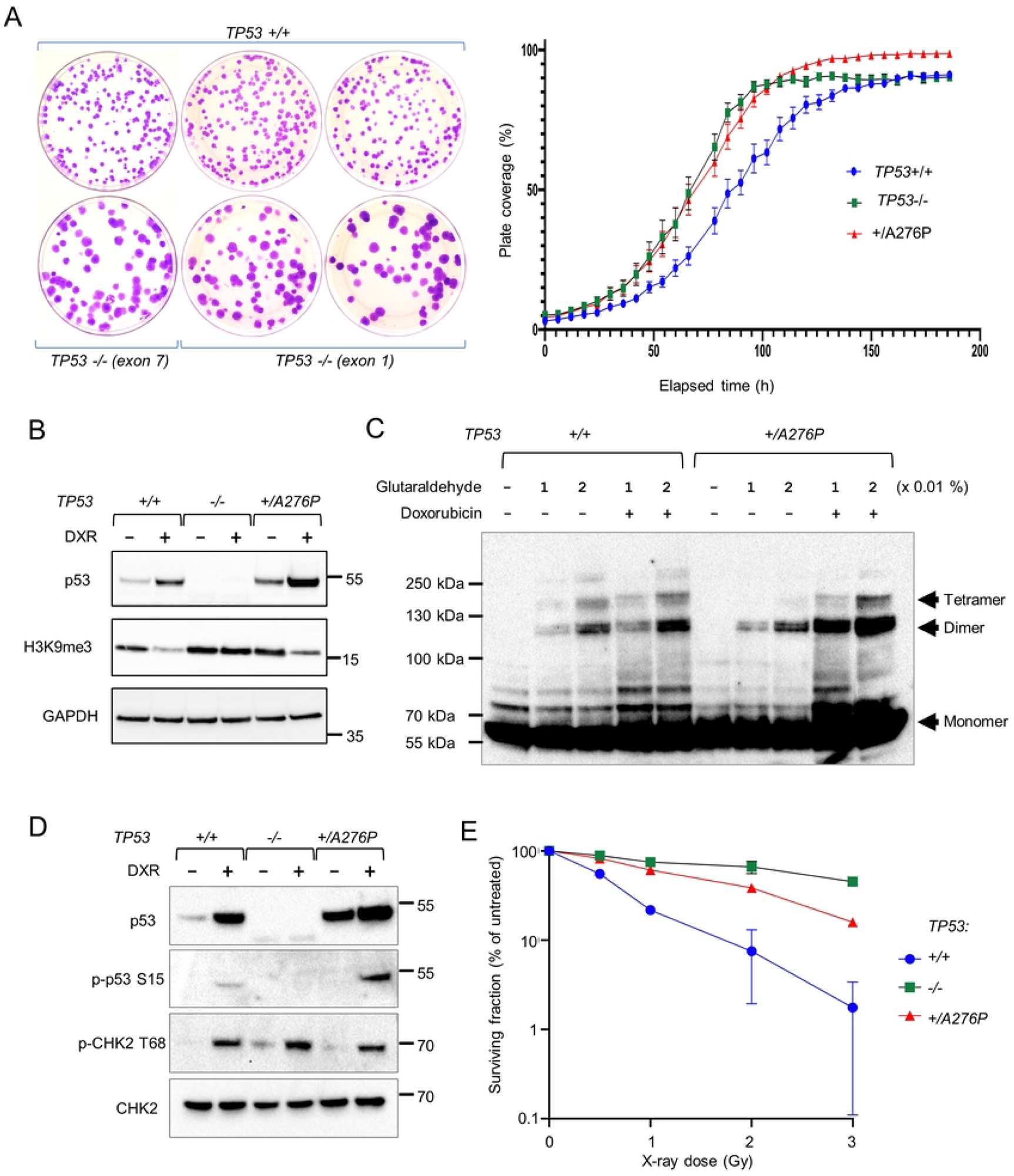
Genetic alteration of *TP53* increased unperturbed cell growth and survival after DNA damage. (A) Approximately 200 cells of the indicated genotypes were plated to 10 cm plates, which were then incubated for 14 d and stained with crystal violet (left) 1.0E+3 cells were plated in a 96-well plate in triplicate wells and placed in an incubator. Cell density was recorded every 6 h by an Incucyte imaging system (right). (B) Monolayer cell cultures were treated with 0.2 µg/ml doxorubicin for 48 h. Extracts were probed for the indicated proteins. (C) Untreated cells and cells treated for 24 h with 0.2 µg/ml doxorubicin were lysed and treated with either 0.01% or 0.02% glutaraldehyde, as indicated. Crosslinked oligomers were detected on a western blot probed with an antibody against p53. (D) Cells were treated with 0.2 µg/ml doxorubicin for 24 h. Whole cell lysates were probed for the indicated proteins and phosphoproteins. (E) Clonogenic survival following irradiation was assessed by counting colonies on three replicate plates per dose, and expressed as a percent of colonies formed by untreated cells of the same genotype.

P53 plays a central role in the cellular responses to DNA damage. Phosphorylated by upstream kinases in the DNA damage signaling network, activated p53 binds to its sequence-specific response elements throughout the genome and induces transcription of numerous downstream genes that collectively restrain growth and promote programmed cell death [17]. The upregulation of transcription by p53 after DNA damage or MDM2 inhibition has been extensively studied, but it is now clear that p53 also silences a subset of its direct targets in the absence of upstream signals [19,20]. The epigenetic repression of transcription by p53 is mediated by the methylation of histone H3K9 [21,22]. A repressive mark, H3K9 trimethylation is continuously maintained in the absence of DNA damage by a chromatin-bound complex containing p53, USP7 and MDM2, which cooperatively recruit the histone methylase SUV39H1 [23]. This complex is rapidly disassembled following DNA damage and the resulting stabilization of p53. With local chromatin in an active euchromatic state, p53 forms DNA-bound tetramers that are required for target gene induction. Conversely, the H3K9me3 mark is elevated in heterochromatin, and maintained in this transcriptionally inactive state by monomeric and dimeric p53. Thus, in a bifunctional manner, p53 can exert tight control over select genes both before and after DNA damage.

We employed the chemotherapeutic agent doxorubicin to stimulate p53 tetramerization and thereby disrupt the complex required for the retention of SUV39H2 at p53 responsive promoters. A global reduction in H3K9me3 protein, previously observed in the colorectal cancer cell line HCT116 [22], is clearly present in *TP53 +/+* RPE1, retained in the *TP53 +/A276P* line but completely absent in p53-deficient RPE1 cells (Fig 2B).

We further examined the effects of DNA damage on p53-dependent gene regulation by assessing oligomeric, chromatin-associated p53 complexes. By crosslinking associated proteins with glutaraldehyde, we were able to resolve endogenous p53 dimers, tetramers and higher order complexes (Fig 2C). In the absence of damage, *TP53 +/A276P* cells exhibited primarily p53 dimers, which are known to be transcriptionally inactive. Cells that expressed only wild type p53, in contrast, exhibited a range of p53 complexes, including transcriptionally active tetramers. Following DNA damage and p53 activation, p53 in wild type cells formed both dimers and tetramers, while the *TP53 +/A549* cells expressed a proportionally lower amount of tetrameric p53, consistent with the reduction in p21 and MDM2 induction in this line (Fig 1D). These patterns suggest that the stable p53 A276P mutant protein primarily formed inactive homo- and heterodimers.

The upstream responses to doxorubicin-mediated DNA damage were intact in each of the cell lines, as evidenced by the retention of their ability to phosphorylate p53 S15 and CHEK2 T68 (Fig 2D), two well-characterized substrates of upstream kinases. To quantify the effects of *TP53* genotype on cell fate after DNA damage, we measured clonogenic survival after exposure to ionizing radiation. The survival of RPE1 cells was strongly reduced by relatively low doses of X-rays (Fig 2E). In contrast, the p53-deficient cells were markedly radioresistant; the *TP53+/A276P* cells exhibited intermediate radiosensitivity, suggesting a wild type gene dosage effect.

The abundance of p53 and its localization are both tightly controlled, primarily by the E3 ubiquitin ligase MDM2. In the absence of DNA damage, p53 largely resides outside the nucleus. In unstimulated cells with low p53 and low MDM2, the nuclear export of p53 is mediated by MDM2-mediated mono-ubiquitination [24]. When MDM2 levels are high, as is the case when cells are recovering from DNA damage, p53 is polyubiquitinated and thus rapidly targeted for degradation by the proteasome. Pulldowns of ubiquitinated p53 revealed increased levels of mono- and polyubiquitination in the *TP53 +/A276P* cells (Fig 3A). The levels of ubiquitinated p53 were similar in the wild type and mutant cells following treatment with the proteasome inhibitor MG-132.

**Figure 3.**
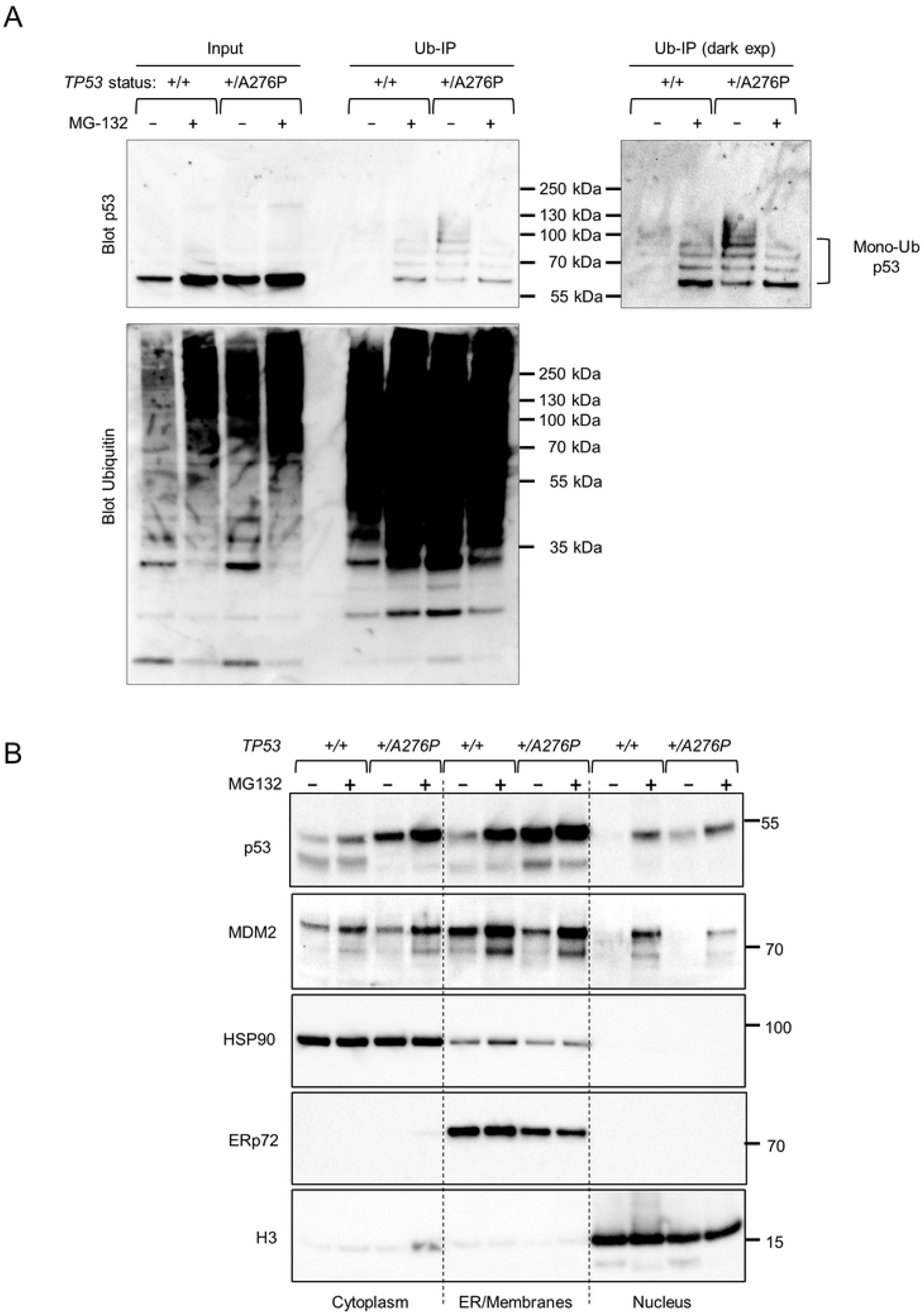
Ubiquitination and nuclear export of wild type p53 and p53 A276P. (A) Untreated cells and cells treated with the proteasome inhibitor MG-132 (10 µM) for 4 h were lysed. Ubiquitinated proteins were pulled down as described in Experimental procedures. Equal amounts of lysate (input) and bead eluate were fractionated and probed for p53 or ubiquitin, as indicated. Two exposures of the same blot are shown to allow visualization of ubiquitinated proteins. (B) Cells treated as in (A) were fractionated into cytoplasmic, intracellular membrane, and nuclear components. Fraction-specific proteins were probed with antibodies against MDM2 and p53. HSP90, ERp72 and histone H3 were detected on a separate blot run in parallel to assess protein recovery.

To investigate the effects of ubiquitination on p53 localization in this system, we fractionated cells to isolate proteins that localized to the cytoplasm or nucleus, or that were associated with the ER and other intracellular membranes (Fig 3B). We found that p53 in the *TP53 +/A276P* cells was increased across all three compartments, but was most abundant in the membrane-associated fraction. The level of MDM2 in this fraction was higher in the wild type cells than in the *TP53 +/A276P* mutant, suggesting that monoubiquitinated wild type p53, exported from the nucleus, was subsequently polyubiquitinated by MDM2 in the membrane compartment. This two-step mechanism would be consistent with the rapid turnover of p53 expressed in wild type RPE1 cells (Fig 1D).

In addition to constitutive ubiquitination by MDM2, phosphorylation by calcium-dependent protein kinase C (PKC) has also been identified as an important requirement for normal p53 turnover in unstressed cells [25,26]. To stimulate p53 turnover via PKC, we treated cells with phorbol 12-myristate 13-acetate (PMA). This phorbol ester is a synthetic analog of diacyl glycerol, the endogenous activator of PKC-mediated signal transduction. *TP53+/+*, *TP53-/-* and *TP53 +/A276P* cells were treated with PMA alone, or with PMA in combination with the MDM2 inhibitor nutlin-3a (Fig 4A). PMA did not appear to affect the stabilization of p53 by nutlin-3a, supporting the current model in which MDM2 and PKC work in concert. Strikingly, PMA administered without nutlin-3a selectively decreased the level of p53 in the *TP53 +/A276P* cell line. The dynamic change in p53 abundance caused by PMA was next assessed over a 12 h period (Fig 4B). By the end of this time course, PMA reduced p53 in both wild type and *TP53 +/A276P* cells to similar, low levels.

**Figure 4.**
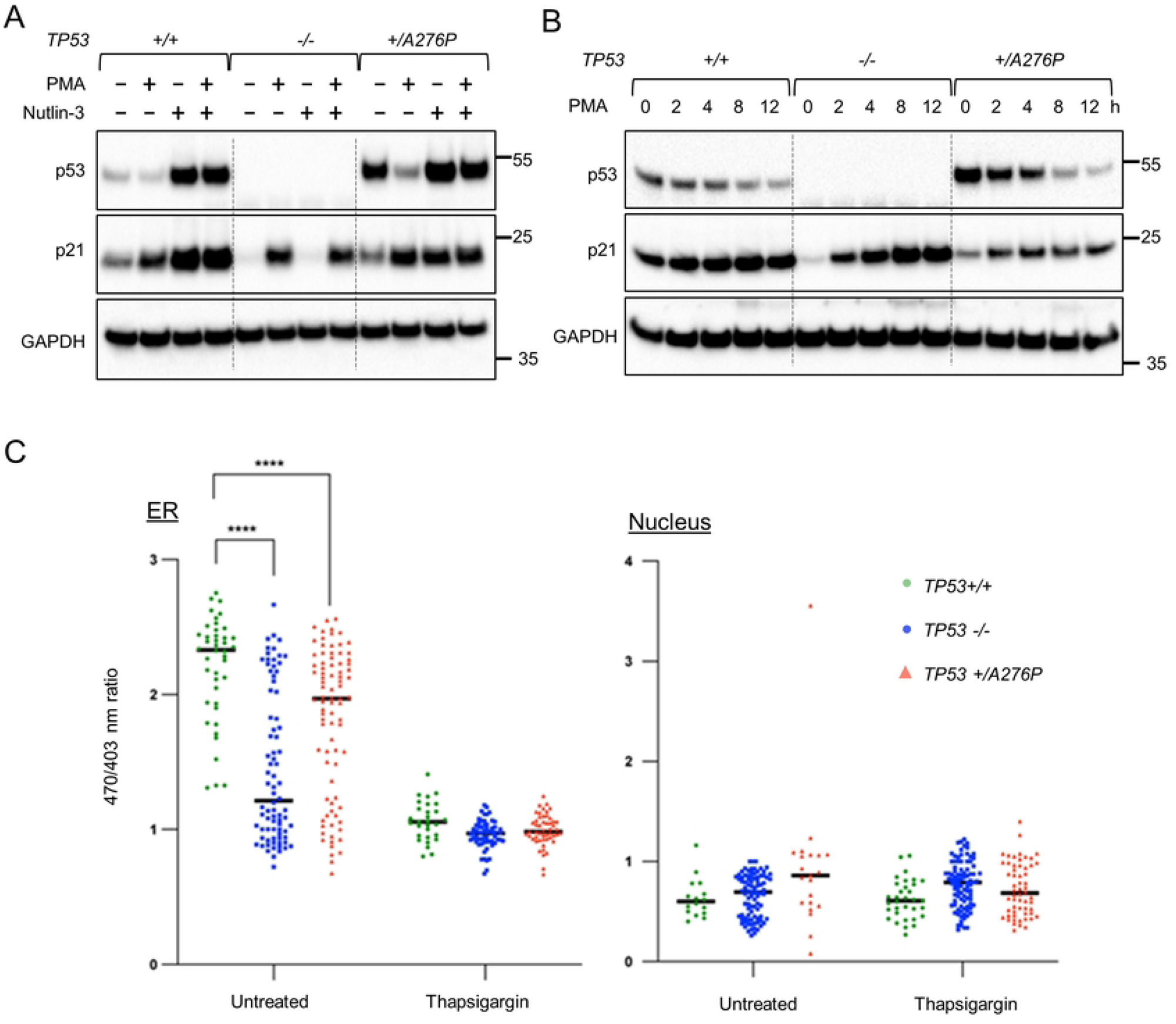
Upstream and downstream effects of p53 on Ca^2+^ signaling. (A) Monolayer cell cultures were treated with nutlin-3a (10 µM) and/or PMA (100 ng/ml) for 8 h. (B) Cells were treated with 100 ng/ml PMA and harvested for protein analysis at the indicated time points. The indicated proteins were assayed by western blot. (C) GAP biosensors were used to assess relative Ca^2+^ levels in the ER and the nucleus. Where indicated, ER Ca^2+^ was selectively depleted by treating cells with the SERCA inhibitor thapsigargin for 24 h.

PMA has antiproliferative effects on some cell lines, which are largely mediated by the p53-independent induction of p21^WAF1/CIP1^ [27–29]. This Ca^2+^ dependent pathway for cell cycle regulation was readily apparent in the RPE1 cell panel (Fig 4A, B). Interestingly, p21^WAF1/CIP1^ upregulation was attenuated in the *TP53 +/A276P* cells. A possible explanation for this observation is that p53 heterodimers in this cell line (Fig 2C) enhanced epigenetic silencing at the *CDKN1A* promoter in the absence of DNA damage. These experiments also illustrate how the turnover of wild type p53 and the A276P mutant are similarly controlled by Ca^2+^-dependent signaling, suggesting a possible therapeutic approach to suppressing the gains-of-function caused by this mutation.

As an important mediator of apoptosis, p53 also plays an upstream role in Ca^2+^ signaling. p53 directly binds to the SarcoEndoplasmic Ca^2+^-ATPase (SERCA) pump at the ER and mitochondria-associated membranes and stimulates the enhanced transfer of Ca^2+^ to the mitochondria [30]. In this transcription-independent manner, wild type p53 increases mitochondrial outer membrane permeability and lowers the threshold for apoptosis.

The abundance of wild type and mutant p53 at the ER-associated membranes (Fig 3B) prompted us to investigate the impact of these proteins on SERCA activity. We used a genetic biosensor to measure the relative levels of Ca^2+^ in the ER lumen, the main repository for intracellular calcium, and in the nucleus. An aequorin-based Ca^2+^ reporter system, insensitive to local pH and Mg^2+^, can be specifically targeted to several organelles via fusion with signaling peptides [31]. Parental RPE1, the p53-deficient knockout and the *TP53 +/A276P* mutant cell line were each stably transfected with reporter constructs encoding a green fluorescent protein (GFP)-Aequorin fusion Protein (GAP) targeted to the ER or to the nucleus. Organelle-specific Ca^2+^ levels were then determined in individual cells by dual-excitation ratiometric imaging, as described [31].

Overall, Ca^2+^ levels in the ER were significantly reduced in *TP53-/-* and, to a lesser extent, in *TP53+/A276P* cells (Fig 4C). This finding is consistent with the recently established role of cytoplasmic p53 as a direct SERCA activator. As the levels of ER-localized p53 were significantly elevated in the *TP53 +/A276P* cell line (Fig 3B), we infer that this mutant protein was non-functional with respect to SERCA activation. In response to SERCA inhibition by thapsigargin, the store of Ca^2+^ in the ER was similarly depleted in each of the three cell lines. The levels of Ca^2+^ in the nucleus were predictably low and did not differ substantially between the three cell lines Fig 4C).

We next examined the effects of the *TP53 +/A276P* mutation on the downstream induction of p53 target genes. By interacting with a bipartite response element in many promoters, wild type p53 directly controls the transcription of genes that collectively suppress cancer phenotypes. The content of the p53-dependent transcriptome varies significantly between cell types and tissues, but commonly includes several canonical targets involved in cell cycle regulation and apoptosis [17,32]. The expression of several well characterized p53-target genes was first assessed by RT-qPCR. The initial panel included *CDKN1A*, which encodes p21^WAF1/CIP1^, *PTCHD4* which encodes PTCH53, *FDXR* which encodes ferredoxin reductase, and *MDM2*. As predicted, each of these target genes was robustly induced in wild type RPE1 following p53 activation by nutlin-3a (Fig 5A). Induction of these genes was notably reduced in the *TP53 +/A276P* cells and minimal or non-detectable in the *TP53 -/-* cells. Particularly for the highly induced genes *CDKN1A* and *PTCH53*, the reduction in gene expression was reduced by more than half in the *TP53 +/A276P* cells, suggesting a dominant negative effect of the A276P mutation on transcriptional transactivation.

**Figure 5.**
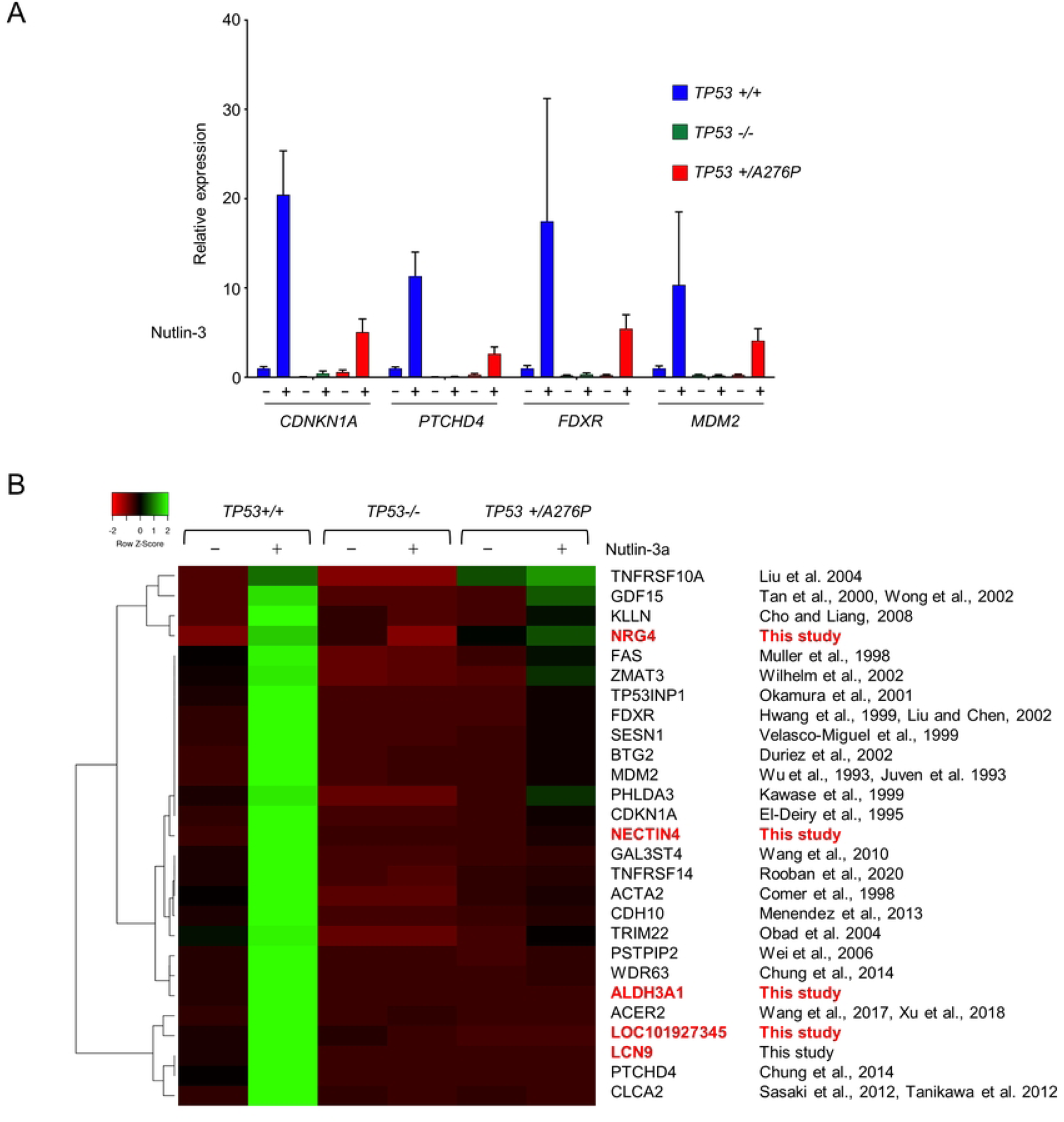
Dominant attenuation of p53-dependent gene expression by p53 A276P. (A) The induction of known p53 target genes by treatment with 10 µM nutlin-3a for 8 h was assessed by RT-qPCR. Expression was normalized to a *GAPDH* control. For each gene the relative expression was calculated in comparison with untreated parental RPE1. (B) RNA-seq analysis of the p53-dependent transcriptome in RPE1. A heatmap illustrates the clustered relationships between genes that were induced at least 3-fold by nutlin-3a in the *TP53+/+* RPE1 cells, and that were also upregulated by at least 10-fold in the nutlin-3a-treated wild type cells over the identically-treated *TP53-/-* cells. Genes previously shown to be p53-regulated are matched with their respective citations at right. Genes highlighted in red were not previously identified as p53 transcriptional targets.

To further assess the impact of the heterozygous *TP53 A276P* mutation on downstream gene expression, we characterized the p53-dependent transcriptome expressed in RPE1 by RNA-seq (Fig 5B). A set of genes that were tightly controlled by p53 in these cells was defined by first identifying those that were induced at least 3-fold in wild type RPE1 after treatment with nutlin-3a for 8 h. This early time point was chosen to eliminate indirect effects caused by upregulation by p53 of other transcription factors. Among these nutlin-3a responsive genes, 27 were induced at least 10-fold higher in wild type RPE1 cells compared with isogenic *TP53-/-* cells that were also treated with nutlin-3a. The induction of this defined transcriptome by nutlin-3a was broadly attenuated in the heterozygous *TP53 +/A276P* cells (Fig 5B).

Interestingly, 22 of the 27 most robustly upregulated genes had previously been linked to p53, with varying levels of confidence, in disparate experimental systems [32–57]. The remaining five genes had not previously been identified as transcriptional targets of p53. However, several of these genes are otherwise linked with p53, cancer or DNA damage. *ALDH3A1* encodes an aldehyde dehydrogenase that was recently shown to confer resistance to oxidative stress by modulating the DNA damage response [58]. *NRG4* encodes a specific ligand for the human epidermal growth factor (EGF) receptor HER4 [59]. *LOC101927345* was recently found to be among a small group of genes that conferred resistance to cisplatin in gastric cancer cells [60]. This heretofore uncharacterized locus encodes the predicted ankyrin repeat-domain-containing protein 20A12. As the ankyrin repeat is a motif that mediates the physical interaction of p53 with several of its binding partners [61], the potential effect of this gene on p53 function may warrant additional investigation.

## Discussion

The telomerase-immortalized cell model described here permits the detailed analysis of p53 phenotypes during unperturbed growth and following exposure to agents that activate p53. We were able to assess transcriptional and non-transcriptional functions of p53 that have been previously characterized individually, in disparate cell types. That these varied phenotypes could be observed and quantified in a single model suggests that the complex pathways upstream and downstream of p53 are intact in RPE1 and highly sensitive to perturbation.

The RPE1 model allowed us to assess the functional impact of a cancer-associated point mutation that arose in one *TP53* allele. Haploinsufficiency caused by the loss of one of the two wild type *TP53* alleles in this cell line as well as dominant negative effects of the heterozygous A276P mutation were readily apparent.

Residue A276 is part of the central DNA binding domain of p53. The nearby residue R273 is among the most highly mutated positions in p53, and one of six recognized mutation hotspots [62]. R273 and A276 form a part of the protein core that directly contacts DNA containing the p53 response element. A previous analysis of amino acid substitutions in the A276 position by Reaz *et al.* [63] revealed that this residue could be replaced by serine (S) or phenylalanine (F) with only subtle effects on promoter selection and transcription. In contrast, our data indicate that the substitution of proline (P) in this position created a mutant p53 protein that was transcriptionally inactive and, moreover, exerted dominant negative effects on transcriptional transactivation by wild type protein. The A276P mutation occurs at low frequency in breast and ovarian cancers, implying that there is selective pressure for expansion of this particular mutation during tumorigenesis. The A276S and A276F mutations created experimentally by Reaz *et al.* are not represented in the TCGA database.

It is unclear how the p53 A276P mutation arose in the RPE1 cell line, which is known to be genetically stable. Codon 276 is located in exon 8 and is 427 bp from the predicted CRISPR cut site in exon 7 (Fig 1A). No other alterations in the region were observed. The Cas9 protein from *S. pyogenes*, employed in our vector, most commonly generates small, localized insertions and deletions (indels), with larger deletions observed at some target sites [64]. Based on the well-described patterns of Cas9-mediated mutagenesis, it would seem improbable that the A276P mutation was an off-target product of gene editing. The expanded cell population was found to be clonally heterozygous, indicating that the mutation must have been present among the cell population from which the subclones were derived. An analysis of short tandem repeats confirmed that this clone was indeed isogenic with RPE1, ruling out the possibility of contamination by cancer cells. For these reasons, we consider it likely that this mutation arose *de novo* during routine cell culture.

It may seem intuitively obvious that driver mutations can arise spontaneously during serial passage of large cell populations, but several studies suggest that this in fact occurs very rarely. Jones *et al.* [65] found that 287 of 289 mutations discovered in human colon cancer xenografts and cancer-derived cell lines were present in the original primary tumor samples. A follow up analysis by Solomon *et al.* of a separate set of samples and a review of the published literature also found no evidence of artifactual genetic alterations caused by *ex vivo* cell culture [66]. Together, these studies should dispel the misconception that there is significant selection for oncogenic mutations and loss of tumor suppressor genes during the serial maintenance of cancer cell cultures. It is tempting to speculate that the apparent suppressive effect of p53 on RPE1 growth, which we have not observed in cancer-derived cellular models, may have allowed this p53 mutation to clonally expand within the parental cell population.

Our rationale for studying p53 in RPE1 was that *in vitro* immortalization by overexpression of telomerase might have spared the p53 pathway from negative selection that would normally occur *in vivo*. Our prediction that these cells would retain particularly robust p53 phenotypes was generally supported by the data reported here. Downstream transcriptional targets of p53 were highly induced after DNA damage or MDM2 inhibition and, conversely, survival was increased in cells that inactivated one or both wild type *TP53* alleles. Interestingly, we found that the effects of p53 were also apparent in the absence of exogenous stimuli. Such phenotypes, which are relatively subtle, have emerged more recently and are an understudied facet of p53 function. In particular, the epigenetic silencing of loci bound by p53 dimers has remained difficult to quantify. Our study suggests that RPE1 cells might be particularly useful for this type of analysis.

We cannot rule out a role for hTERT overexpression in the robust DNA damage phenotypes we observed in RPE1. Essential for the immortal growth of this cell line, telomerase activity is known to augment the cellular responses to exogenous DNA damage, possibly by capping chromosome ends and thereby reducing endogenous DNA damage [67]. Knockdown or heterozygous knockout of hTERT has been shown to cause DNA damage and increase radiosensitivity [68,69]; it is unknown if hTERT overexpression might have opposite effects.

Irrespective of the possible impact of hTERT overexpression, parental RPE1 cells were sensitive to clinically-relevant doses of ionizing radiation while the p53-deficient derivative line was radioresistant (Fig 2E). A causal association between the mutational loss of p53 and radioresistance has been supported by clinical studies [70–72] and by studies of preclinical animal models [73,74]. In contrast, studies of the relationship between p53 status and resistance to ionizing radiation performed in diverse cancer cell line panels have been considerably less definitive [70,75]. While a sensitizing effect of p53 has recently been observed in several isogenic systems generated by gene editing [76,77], such effects have been consistently absent in the original HCT116 isogenic cell pair, whether assayed *in vitro* [68] or *in vivo* [5].

The suppression of RPE1 growth by p53 was likely mediated by the cyclin-dependent kinase inhibitor p21^WAF1/CIP1^, which was expressed at a higher basal level in wild type cells compared with the isogenic p53 knockout line (Figs 1D and 5A). The detection of chromatin-associated p53 tetramers in the absence of exogenous DNA damage (Fig 2D) further supports an active role for p53 in the suppression of unperturbed cell growth in this model. Notably, p53-deficiency in HCT116 does relieve growth inhibition, but this phenotype is dependent on an increase in angiogenesis and is therefore only expressed in cell-derived xenografts [78].

In their seminal study of cytoplasmic p53 and its effects on Ca^2+^ flux, Giorgi *et al.* focused on the non-transcriptional effects of p53 in response to apoptotic stimuli [30]. In our system, a controlling effect of p53 on Ca^2+^ homeostasis was readily apparent in the absence of exogenous stressors. Notably, the study Giorgi *et al.* showed that the ability of p53 to control Ca^2+^ flux is a function that is lost in the hotspot p53 mutants R175H and R273H. Our results suggest that p53 A276P, a mutant found in far fewer cancers, is similarly defective.

Interestingly, there appeared to be a bimodal distribution of Ca^2+^ content among the p53-knockout and, to a somewhat lesser extent, in the *TP53 +/A276* cell populations (Fig 4C). The basis for this interesting distribution is unclear. During its catalytic cycle, SERCA is known to function in two distinct structural and biochemical states that differ in their affinity for Ca^2+^ [79]. It is possible that the interaction between SERCA and p53 favors the transition between these two states, which is less efficient in the absence of p53. Alternatively, this bimodal distribution may simply reflect the dynamic changes in intracellular calcium that are known to occur during the cell division cycle, in which case the effect of p53 on SERCA would be indirect. Additional investigation will be needed to determine how p53 impacts SERCA at the single cell level.

The maintenance of tissue homeostasis by the p53 pathway involves the concerted activities of numerous upstream regulators and downstream effectors. Arguably, the full complexity of this expansive signaling network is unlikely to be captured by any single model system. Complementary studies of different in vitro and in vivo models, including the immortalized system described here, will undoubtably be needed to form a more complete picture of p53-mediated tumor suppression.

## Materials and Methods

### Cell lines and cell culture

A puromycin-sensitive derivative of hTERT-RPE1 was a gift from Andrew Holland. Cells were routinely grown at 37°C in 5% CO2 in DMEM/F12 supplemented with 6% fetal bovine serum (FBS) and penicillin/streptomycin. The parental cell line and all derivatives were authenticated by Short Tandem Repeat (STR) profiling and tested for the presence of mycoplasma at the Johns Hopkins Genomic Resources Core Facility.

### Generation of TP53-/- cells

For the disruption of TP53 exon 7, an oligonucleotide duplex encoding the CRISPR guide sequence 5-GCATGGGCGGCATGAACCGG-3 was directly cloned into the plasmid vector pSpCas9(BB)-2A-Puro (PX459) V2.0, a gift from Feng Zhang (Addgene #62988). The resulting plasmid was introduced into RPE1 by transfection with Lipofectamine 3000 (ThermoFisher Scientific). Following a 4 d selection in 2 µg/ml puromycin, the remaining cells were plated to limiting dilution in 96-well plates. Individual subclones were expanded and screened by PCR, using the forward primer 5’-CTCCTAGGTTGGCTCTGACTGT-3’ and the reverse primer 5-AAACTGAGTGGGAGCAGTAAGG-3. Genetic disruption of both alleles was assessed by Sanger sequencing followed by analysis with Inference of CRISPR Edits software (Synthego). The deletion of *TP53* exon 1 was accomplished by a similar approach. The flanking guides 5’-TAGTATCTACGGCACCAGGT-3’ and 5’-TCAGCTCGGGAAAATCGCTG-3’ were designed to create a 385 bp deletion that included the entire exon. The expected deletion was identified in multiple subclones by PCR with the forward primer 5’-CTCCAAAATGATTTCCACCAAT-3’ and the reverse primer 5’-ACTTTGAGTTCGGATGGTCCTA-3’. For all knockout clones, the loss of p53 expression was confirmed by western blot.

### Identification of the TP53 A276P mutation

Each of *TP53* exons in the clone that overexpressed p53 was amplified by PCR and sequenced. A single mutation was identified in exon 8, which was amplified by the forward primer 5’-CTTAGGCTCCAGAAAGGACAAG-3’ and the reverse primer 5’-AGAGGCAAGGAAAGGTGATAAA-3’

### Western blots, antibodies and cell fractionation

Protein lysates were prepared in RIPA buffer (Cell Signal Technologies), resolved on Bolt Bis-Tris minigels (ThermoFisher Scientific) and transferred to PVDF membranes (MilliporeSigma). Antibodies for the detection of p53 (DO-1) and MDM2 (SMP14) were obtained from Santa Cruz Biotechnology. Antibodies recognizing p21^WAF1/CIP1^ (12D1) and H3K9me3 (D4W1U) phospho-Chk2 (T68, polyclonal), phospho-p53 (S15, 16G8), HSP90 (C45G5), ERp72 (D70D12) and histone H3 (D1H2) were purchased from Cell Signaling Technology. The isolation of ubiquitinated proteins was performed with the Signal-Seeker Ubiquitin Enrichment kit (Cytoskeleton). Cell fractionation was performed with the Qproteome Cell Compartment kit (Qiagen).

### Glutaraldehyde crosslinking

Multimeric forms of p53 were stabilized by glutaraldehyde crosslinking, as previously described [80]. Briefly, cells were lysed with a buffer containing 0.5% NP-40 substitute. Lysates were brought to a final concentration of glutaraldehyde of 0.01 or 0.02% and incubated on ice for 20 min. The crosslinking reaction was stopped with 1X Bolt sample buffer (ThermoFisher Scientific) and proteins were resolved by gel electrophoresis followed by a western blot, as described above.

### Drug treatments

The MDM2 inhibitor nutlin-3a and the proteasome inhibitor MG-132 were both purchased from Enzo Life Sciences, dissolved in DMSO and used at final concentrations of 10 µM and 20 µM respectively. The SERCA inhibitor thapsigargin and doxorubicin were purchased from Cell Signaling Technology and dissolved in DMSO. Cycloheximide, used for the assessment of protein stability, was purchased as a ready-made solution (Sigma Aldrich) and used at a final concentration of 100 µg/ml. Phorbol-12-myristate-13-acetate (PMA) was also obtained from Sigma Aldrich, dissolved in ethanol, and used at a final concentration of 100 ng/ml.

### Clonogenic survival assay

For the assessment of clonogenic survival following irradiation, between 200-500 cells were plated to 10 cm cell culture dishes in triplicate. After allowing cells to attach for 16-20 h, plates were exposed to measured doses of X-rays delivered with a MuliRad225 (Faxitron) and returned to the incubator for 14 d. Colonies were fixed and stained with 0.2% crystal violet in methanol, and colonies containing >50 cells were counted on a plate scanner (Interscience).

### Imaging of subcellular Ca^2+^

Cells were transfected with the plasmid pcDNA3_erGAP2 (a gift from Teresa Alonso and Javier García-Sancho, Addgene #78120), which encodes a low affinity fluorescent calcium biosensor, based on a GFP-aequorin fusion protein (GAP), targeted to the endoplasmic reticulum. A similar plasmid, pcDNA3_nucGAp (a gift from Teresa Alonso, Addgene #78736) was used to assess nuclear Ca^2+^. Dual-excitation imaging of GAP-expressing cells were performed on a fluorescent microscope equipped with 403 and 470 nm excitation filters.

### Analysis of gene expression

The induction of p53 target genes was assessed by quantitative reverse transcription PCR (RT-qPCR). Total RNA was extracted from subconfluent cells with the Monarch total RNA purification kit (NEB), and reverse transcribed with LunaScript RT SuperMix (NEB). Real time PCR amplification was performed on a BioRad CFX96 Real-Time PCR detection system using the following primer sets: *CDKN1A* (p21^WAF1/CIP1^): forward 5’AGGTGGACCTGGAGACTCTCAG-3’, reverse 5’-TCCTCTTGGAGAAGATCAGCCG-3’; *PTCHD4* (PTCH53): forward 5’-AAGCCAGCTCTATTCGGACTTAC-3’, reverse 5’-ACTTGACCCGCTGATCTTTG-3’; *FDXR*: forward 5’-TCTTATACCCAATGCTGCTGAG-3’, reverse 5’-TCACTAGACTGGAGGGTGTC-3’; *MDM2*: forward 5’-GAGAGCAATTAGTGAGACAGAAGA-3’, reverse 5’-GCTTTCATCAAAGGAAAGGGAAA-3’.

### RNA-seq analysis

Total RNA was extracted from each cell line, either untreated or following treatment with 10 µM nutlin-3a. Poly(A) selection, library preparation and 2×150 bp Illumina sequencing were performed by Azenta Life Sciences. A total of 462,496,793 reads encompassing 138,751 Mb were obtained from the six samples (single replicates). Reads were imported into Geneious Prime (Version 2020.0) for processing. Reads were first paired and trimmed with BBDuk, then mapped to Hg38 with the Geneious RNA Mapper. For the calculation of normalized expression levels in transcripts per million (TPM), ambiguously mapped reads were counted as partial matches. Differential expression between samples was determined in a pairwise fashion using the median of gene expression ratios, as described [81]. The normalized read counts across samples were used to generate z-scores for each row. Genes that met the specified criteria were clustered by the complete linkage method using the Heatmapper online application (http://heatmapper.ca/expression/). Distance between rows was measured by the Spearman Rank Correlation.

## Acknowledgements

We thank the many colleagues who generously provided reagents that were used in this study. The results here are in part based upon data generated by the TCGA Research Network: https://www.cancer.gov/tcga. Sequence data generated in this study have been deposited at the Gene Expression Omnibus (GEO) DataSet repository, accession number GSE229160.

## Abbreviations and nomenclature

5-FU: 5-fluorouracil
PKC: calcium-dependent protein kinase C
PMA: phorbol 12-myristate 13-acetate
SERCA: SarcoEndoplasmic Ca^2+^-ATPase

